# NIHBA: A Network Interdiction Approach with Hybrid Benders Algorithm for Strain Design

**DOI:** 10.1101/752923

**Authors:** Shouyong Jiang, Yong Wang, Marcus Kaiser, Natalio Krasnogor

## Abstract

Flux balance analysis (FBA) based bilevel optimisation has been a great success in redesigning metabolic networks for biochemical overproduction. To date, many computational approaches have been developed to solve the resulting bilevel optimisation problems. However, most of them are of limited use due to biased optimality principle, poor scalability with the size of metabolic networks, potential numeric issues, or low quantity of design solutions in a single run. In this work, we have employed a network interdiction model free of growth optimality assumptions, a special case of bilevel optimisation, for computational strain design and have developed a hybrid Benders algorithm (HBA) that deals with complicating binary variables in the model, thereby achieving high efficiency without numeric issues in search of best design strategies. More importantly, HBA can list solutions that meet users’ production requirements during the search, making it possible to obtain numerous design strategies at a small runtime overhead (typically *∼*1 hour).

## 1 Introduction

With the advance of genome-scale metabolic modelling (GSMM), the past decades have witnessed a significant number of computational tools for microbial metabolic engineering [Maia *et al.*, 2016]. These tools facilitate improved strain performance for the production of a variety of high-value biochemicals and biosynthetic precursors, including vanillin [Brochado *et al.*, 2010], lycopene [Choi *et al.*, 2010], malonyl-CoA [Xu *et al.*, 2011], alkane and alcohol [Fatma *et al.*, 2018].

A large number of strain design tools are based on bilevel optimisation. OptKnock [Burgard *et al.*, 2003] is one of the earliest bilevel optimisation based tools. OptKnock maximises target chemical production while assuming mutant strains at optimal growth in flux balance analysis. The resulting bilevel problem is solved through a reformulation that makes the inner level problem equivalent constraints under the condition of strong duality [Burgard *et al.*, 2003]. The OptKnock model was latter extended to improve target production via gene up/down-regulation [Pharkya and Maranas, 2006], cofactor specificity [King and Feist, 2014], or heterologous pathways [Pharkya *et al.*, 2004], and to develop anti-cancel drugs by the identification of synthetic lethal genes [Pratapa *et al.*, 2015]. These studies demonstrate the great effectiveness of the bilevel optimisation based framework in metabolic engineering.

However, the bilevel optimisation based framework in literature has numerous limitations. The first one is the intensive computational cost in search of optimal solutions. Bilevel optimisation is often reformulated into a mixed-integer linear programming (MILP) so as to be solved by exact MILP solvers. It can take up to a week to solve an MILP resulting from a medium-sized GSMM [Feist *et al.*, 2010]. Many practical strategies, such as model reduction and refinement of candidate knockout set [Feist *et al.*, 2010], have been used to reduce the computational time but may miss the best design strategies due to reduced search space. GDBB introduced a truncated branch and bound to speed up the search process. GDLS used local search with multiple search paths to reduce the search space for each local MILP [Lun *et al.*, 2009]. While finding optimal solutions are computationally costly for exact solvers, other studies resorts to inexact methods, such as genetic algorithms [Patil *et al.*, 2005; Rocha *et al.*, 2010] and swarm intelligence [Choon *et al.*, 2015]. These methods, however, still scale poorly with the size of GSMM and are specially ineffective when a large number of genetic manipulations are allowed for high target production.

In company with intensive computations, the resulting MILP often has weak LP relaxations due to disjunctive big-M constraints [Codato and Fischetti, 2006], another limitation of the current bilevel optimisation based methods. Big-M formulation can easily cause numeric issues, particularly in genome-scale metabolic models where stoichiometric coefficients often vary many orders of magnitude [Sun *et al.*, 2013]. As a result, optimal strain design solutions returned from global search solvers such as Gurobi and CPLEX may turn out to be *in silico* infeasible. Model reformulation may alleviate numeric issues and potentially reduce computational costs. However, a proper model reformulation is often time-consuming and laborious as extra care has to be taken to prevent other numeric difficulties while fixing one.

The third limitation is that only a single solution is obtained in each execution of optimisation by modern solvers. Multiple runs are required to generate more solutions, which inevitably increases computational burden. Despite that some commercial solvers, such as Gurobi and CPLEX, provide options to preserve multiple solutions in a single run, many alternative solutions exist for the optimal production rate in underdetermined metabolic systems, and they are likely to have similar production envelopes [Lewis *et al.*, 2012]. This similarity renders the bilevel optimisation less attractive as little information can be gained about the trade-off between growth and production for decision making. Also, these exact solvers consider two solutions different when their continuous but integer variables have different values. Such solutions lead to same design strategies, which is of not interest to decision makers. Heuristic methods, such as local search in GDLS [Lun *et al.*, 2009] and population-based algorithms [Jiang *et al.*, 2018; Patil *et al.*, 2005] may help to find diverse solutions but often suffer from local optima.

Another limitation in most of bilevel optimisation based tools is potential biases induced by the optimisation principle in the inner-level FBA [Lewis *et al.*, 2012]. OptKnock and many of its derivatives assume mutant strains have a biologically meaningful objective which is often to maximise growth. However, this assumption is not always correct as some microorganisms seem to achieve a multiobjective tradeoff of metabolism [Schuetz *et al.*, 2012], and mutants prefer small metabolic adjustments from the wild type [Segré *et al.*, 2002]. It is there desirable that bilevel models eliminate the biased assumption on cell growth while optimising the target production rate.

There are also a number of tools free of bilevel optimisation. For example, approaches based on minimum cut set (MCS) identification have also developed to remove all possible design strategies that do not meet specific requirements of a desired production strain[Apaolaza *et al.*, 2018; von Kamp and Klamt, 2014]. MCS based approaches, however, are also computationally intensive as they need to enumerate elementary modes of a given metabolic network. OptForce identifies genetic interventions by investigating the difference in flux distributions between the wild type and the desired mutant [Ranganathan *et al.*, 2010]. OptForce showed good predictions in *in vivo* studies [Xu *et al.*, 2011]. However, the requirement on flux measurements of the wild type, which is not always available, limits its wide applicability.

In view of these limitations in existing bilevel optimisation based tools, this paper considers the identification of genetic manipulations as network interdiction (NI), a problem well studied in game theory [Lim and Smith, 2007]. The NI involves one game player (host strain) that tries to avoid the overproduction of target chemicals for cellular homeostasis whereas the other opposing player (metabolic engineer) attempts to manipulate the metabolic network in order to maximally disrupt the first player’s activity. Therefore, the NI is a max-min problem in which the objective involves only the target production, avoiding the use of the widely assumed growth optimality. The NI is a special case of general bilevel problems. The solution to this NI problem is a novel hybrid algorithm based on Benders decomposition [Codato and Fischetti, 2006], aiming to address the other limitations mentioned previously. NIHBA, the proposed approach, has shown the ability to efficiently find a large number of growth-coupled design strategies with diverse production envelopes in a single run, and to scale well with the size of allowable knockouts.

## 2 Results

### 2.1 NIHBA: using network interdiction and Benders decomposition

More often than not, wild type strains maintain homeostasis and thus avoid overproducing a product of interest while maximising biomass (Fig. 1A). Metabolic engineering of host strains requires metabolic network modifications leading to improved flux towards the biosynthetic pathway of the target product. Metabolic engineers who look for the best modifications can be considered adversaries or interdictors as they use limited engineering costs (time, financial cost, etc.) to intervene the host’s activities and reverse the cell’s low target production (Fig. 1A). This strain design task is similar to network interdiction (NI), a game theoretic framework where a budget-constrained interdictor intervenes a network user’s activity, e.g. commodity flow [Lim and Smith, 2007], by removing network arcs.

**Figure 1:**
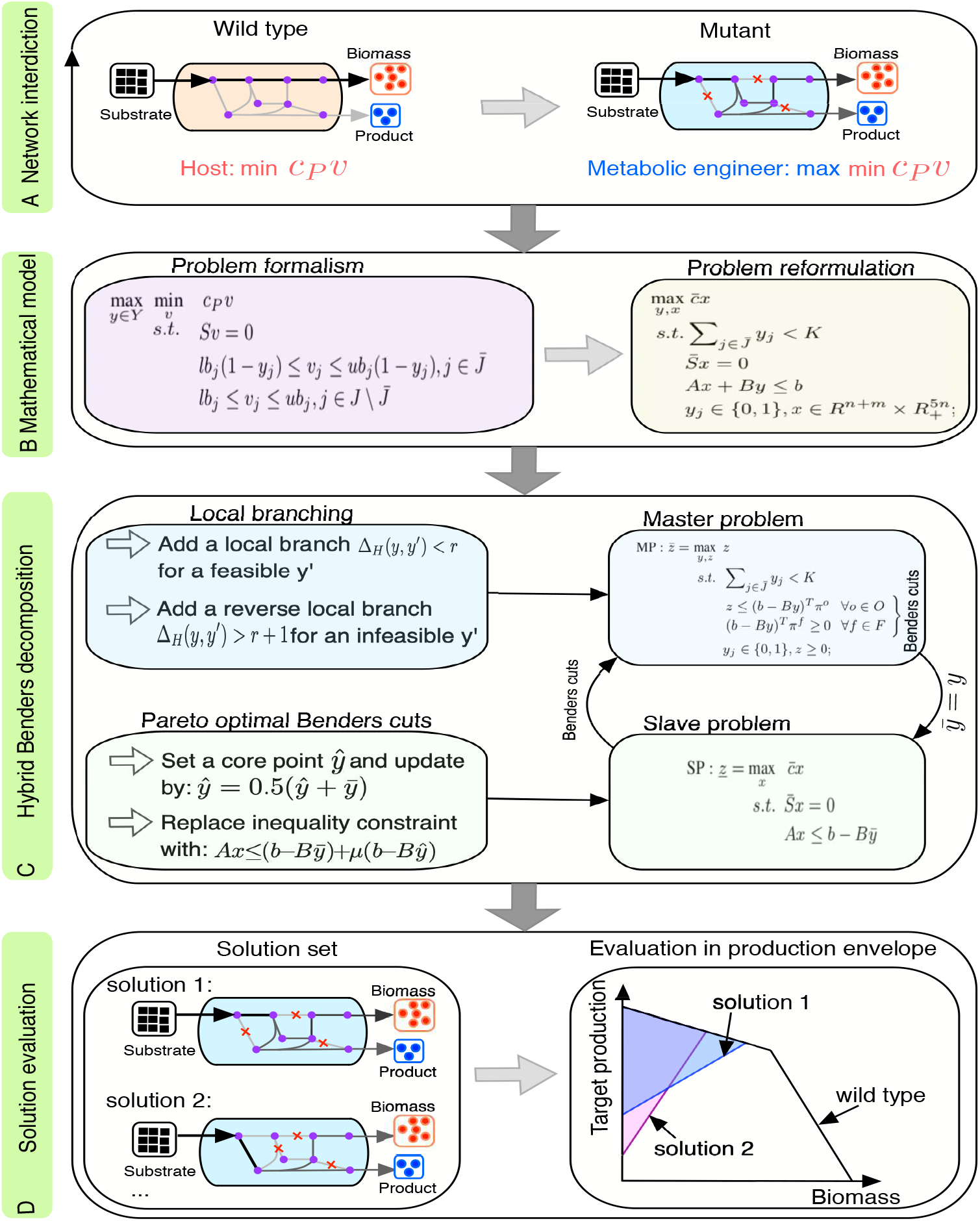
A schematic workflow of the proposed NIHBA tool for strain design. (**A**) Illustration of network interdiction in strain design: host cells avoid overproducing a product whereas metabolic engineers interdict the host network to maximally impair the host’s activity. (**B**) Mathematically modelling the network interdiction problem in strain design, followed by problem reformulation to obtain a standard MILP problem. (**C**) Hybrid Benders decomposition algorithm. The MILP is decomposed into a binary master problem and a linear slave problem, and Pareto-optimal cut generation and local branching are introduced to speed up the search of solutions. (**D**) Solutions that meet production requirements are stored and evaluated in production envelopes.

We proposed a NI model for identifying gene-associated reaction knockouts, but up/down regulation of genes can be considered in this model as well. The NI model is a special case of bilevel optimisation. It was recast into a standard MILP problem (Fig. 1B) using a special reformulation approach (Material and Methods). The resulting MILP contains both complicating binary variables and easy continuous variables. It can be computationally intensive for a large size of binary variables and/or a high allowable number of knockouts, and likely to have numeric issues for global search due to Big-M effects [Codato and Fischetti, 2006]. We therefore resorted to Benders decomposition for this NI model. We proposed a hybrid Benders algorithm (HBA) with two novel techniques to solve the model efficiently and obtain as many design solutions as possible in a single run (Fig. 1C). The solutions from our approach, NIHBA, were then analysed in production envelopes (Fig. 1D), from which the most promising design solution can be selected for implementation.

### 2.2 Case studies

NIHBA was tested on iML1515 [Monk *et al.*, 2017], the largest GEM model for E. coli, for the production of succinate and ethanol. For large models, Feist *et al.* [2010] suggested a preprocessing procedure to ease computational costs. In particular, reactions involving compounds that have more than a certain number (*n*_*c*_) of carbons are not considered as knockout candidates since they are assumed unlikely to carry high flux. We tested different values of *n*_*c*_ = {10, 15, 22, 100} (*n*_*c*_ = 100 indicates no reaction removed due to carbons), resulting in different sizes of candidate set *n*_*s*_ = {152, 204, 272, 342}. In all simulations, designed strains were required to have at least 10% cell growth of the wild type.

We show in Table 1 that the reduction of candidate set by carbon number has a significant effect on target production, especially for succinate. Exclusion of reactions with a carbon number of over 10 (corresponding to 152 candidate knockouts) results in a low succinate production flux of 6.8838 *∼* 9.9430 mmol/gDW/h, which is less than a third of the theoretic maximum production (TMP). A slight relaxation of carbon number to 15 helps to identify a solution with around two thirds of TMP, and succinate production reaches *∼*25 mmol/gDW/h(73% TMP) when no carbon number is constrained in candidate reactions. This indicates that some actions with large carbon numbers are very important in redirecting flux towards succinate, although they may not carry high flux values. For example, both reactions PDH and PFL acting on a high carbon compound, i.e., Acetyl-CoA, carry a good flux value in wild type strains. The knockout of them together with TALA and LDH L under anaerobic condition predicts high succinate production by our method. This prediction agrees well with *in vivo* studies [Hong and Lee, 2002; Zhu *et al.*, 2007]. It is also observed that many solutions are found by NIHBA, and all of them renders a growth-coupled production phenotype. Particularly, there exist a vast number of growth-coupled solutions with low succinate production rate, as indicated when the size of the candidate knockout set is *n*_*s*_ = 152. Most of them do not appear in larger candidate sets, implying that NIHBA prefers high-production solutions. In the following, a set of 342 candidate reactions is used for NIHBA.

**Table 1:** Succinate and ethanol production predicted by NIHBA with at most five knockouts for different sizes of candidate set. The growth rate, minimum production and maximum production are from the solution with the highest minimum production rate.

| $n_s$ | succinate | | | | ethanol | | | |
| --- | --- | --- | --- | --- | --- | --- | --- | --- |
|  | #solutions | growth<br>(h<sup>-1</sup>) | min. prod.<br>(mmol/gDW/h) | max. prod.<br>(mmol/gDW/h) | #solutions | growth<br>(h<sup>-1</sup>) | min. prod.<br>(mmol/gDW/h) | max. prod.<br>(mmol/gDW/h) |
| 152 | 1255 | 0.8729 | 6.8838 | 9.9430 | 21 | 0.1238 | 38.0668 | 38.0986 |
| 204 | 186 | 0.2023 | 21.1032 | 24.3682 | 37 | 0.1238 | 38.0668 | 38.0986 |
| 272 | 271 | 0.1908 | 22.2077 | 25.0055 | 52 | 0.1238 | 38.0668 | 38.0986 |
| 342 | 172 | 0.1549 | 25.0252 | 25.0660 | 84 | 0.1187 | 38.1716 | 38.2466 |

In contrast to succinate, ethanol can be easily produced at a high rate (e.g., 38mmol/gDW/h, or equivalently 95% TMP) by knocking down only reactions of low carbon number. Therefore, excluding as many reactions of high carbon number is beneficial for computational efficiency while achieving a high-production design strategy. The number of growth-coupled solutions for ethanol is, however, much fewer than that for succinate.

Next, we analyse the performance of NIHBA on a varying number of knockouts. We observe that, for both succinate and ethanol, the production rate increases sharply when more knockouts are allowed but levels out from five knockouts (Fig.2A and B). The number of growth-coupled solutions and the percentage of high-production solutions increase with the allowable number of knockouts. No solutions with >80% TMP were found within 15 knockouts for succinate, and only a tiny portion of solutions has a production rate of <20% TMP for ethanol. This indicates again that NIHBA favours high-production solutions during the search.

**Figure 2:**
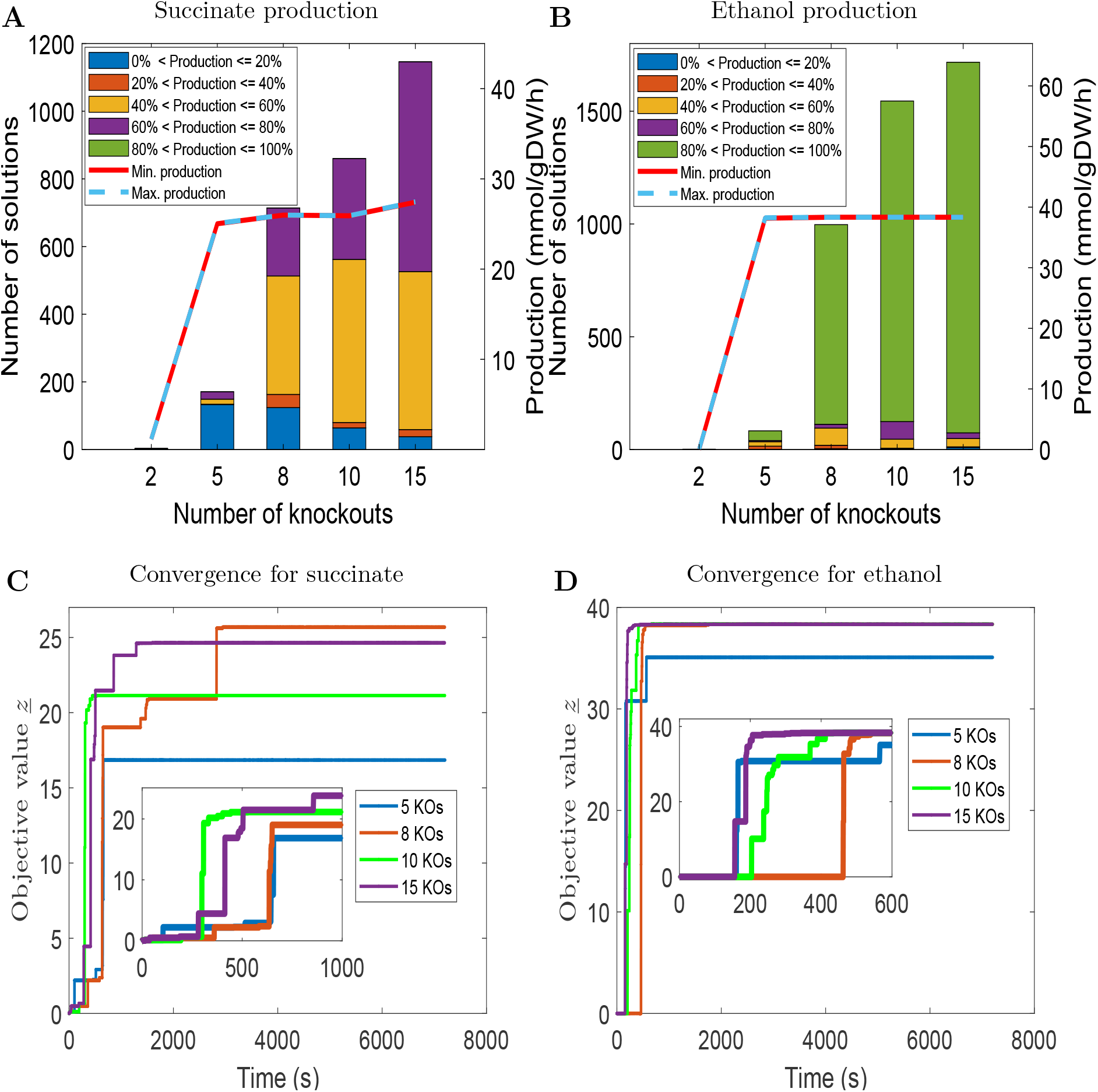
Performance of NIHBA with different number of knockouts for succinate and ethanol production. (**A**) For succinate, the number of solutions achieving a certain percentage of the maximum theoretic production rate, and the minimum and maximum production rate of the best solution found for a different number of knockouts. (**B**) For ethanol, the number of solutions achieving a certain percentage of the maximum theoretic production rate, and the minimum and maximum production rate of the best solution found for a different number of knockouts (KOs). (**C**) The objective value of NIHBA against runtime for succinate production. (**D**) The objective value of NIHBA against runtime for ethanol production.

We show in Fig. 3 the distribution of single knockouts of obtained solutions in different subsystems and their frequency in design solutions to understand which subsystems/knockouts are likely to be engineering targets. Specifically, all the solutions with >60% TMP identified from a limit of 8 knockouts were analysed in terms of knockout occurrence in different subsystems. We find that extracellular exchange reactions (mainly oxygen uptake), reactions from glycolysis, gluconeogenesis, pyruvate metabolism and pentose phosphate pathway are most likely knockout targets for the production of succinate and ethanol, which shows a good agreement with existing studies [Feist *et al.*, 2010; Tepper and Shlomi, 2010]. All the solutions suggest oxygen depletion (Fig. 3C), which is consistent with the common sense that the two products are best produced in anaerobic environments. It is observed that glucose-6-phosphate isomerase (PGI) appears frequently in design solutions for both products, and high succinate-producing strategies additionally favour ATP synthase (ATPS4rpp) (Fig. 3C). Similar results have been also reported in a recent study [Dinh *et al.*, 2018], where computationally intensive strain design tools were used to obtain a large collection of knockout designs.

**Figure 3:**
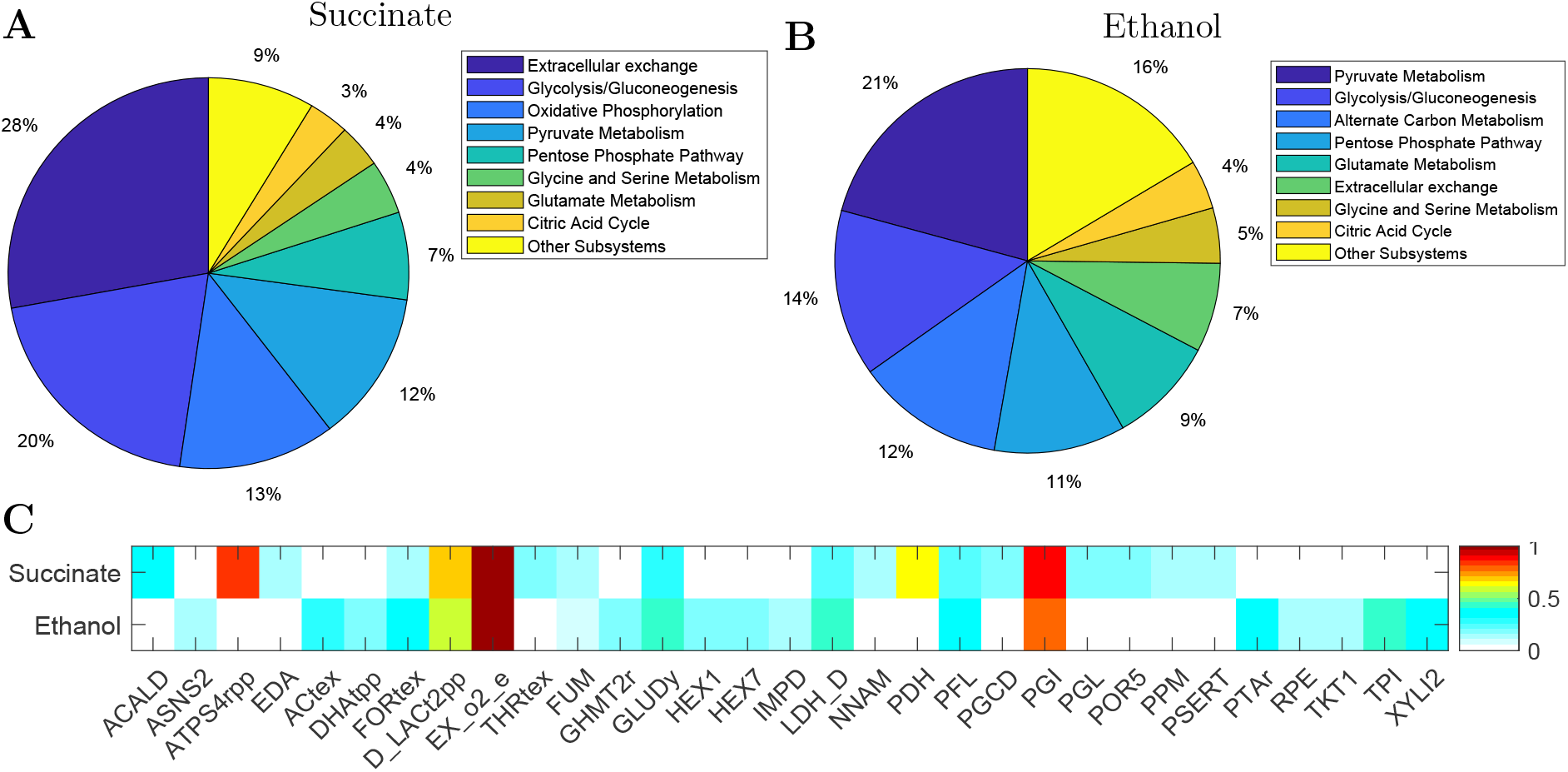
Knockout distribution for succinate and ethanol production. (**A**) The percentage of solutions with at least one knockout from each subsystem for succinate production. (**B**) The percentage of solutions with at least one knockout from each subsystem for ethanol production. (**C**) The fraction of design solutions that have a specific knockout (only knockouts that appear in ¿10% design solutions for either target products are displayed).

The computational cost is low for NIHBA, as shown in Fig.2C and D. A short runtime (300*∼*600s) enables NIHBA to identify high-production solutions for both succinate and ethanol, indicating that our hybrid approach can quickly generate effective Benders cuts to reduce search space. Depending on the maximum allowable number of knockouts, the runtime required before reaching the convergence stage is different, but with a small variation. Generally, NIHBA starts to converge after *∼*2000s and 600s for succinate and ethanol, respectively. The longer time required for succinate may be explained by relatively fewer high-production solutions in the design space. Despite that, the computational time required by NIHBA for a good solution is small (compared to days*∼*weeks in existing methods [Feist *et al.*, 2010]) and does not increase exponentially with the number of knockouts, a widely recognised issue in global search [Feist *et al.*, 2010; Lun *et al.*, 2009].

### 2.3 Comparison with other tools

For comparison, the network interdiction problem was also solved using the OptKnock [Burgard *et al.*, 2003] and GDLS [Lun *et al.*, 2009] approaches with Gurobi [Gurobi Optimization, 2018], called NI-OptKnock and NI-GDLS, respectively. NI-GDLS used *M* = 5 search paths and a search size of *k* = 3 in order to get multiple solutions. For efficiency, parameters in the Gurobi solver was set according to Egen and Lun [2012].

For succinate, when at most five knockouts are allowed, NIHBA found a large number of solutions whereas both NI-OptKnock and NI-GDLS failed to find a feasible solution. The failure is mainly due to numeric issues in Big-M formulation, which existed even although we switched to indicator constraints or CPLEX12.8 for MILP. This demonstrates that HBA overcomes such numeric issues. For readability, we only show a small number of selected solutions from NIHBA in the production envelope (Fig. 4A). As seen, NIHBA can obtain diverse solutions forming a good representative of the tradeoff between cell growth and succinate production. Interestingly, NIHBA identified a strong growth coupled design (non-zero production at no growth), despite its slightly suboptimal production rate at the maximum growth.

**Figure 4:**
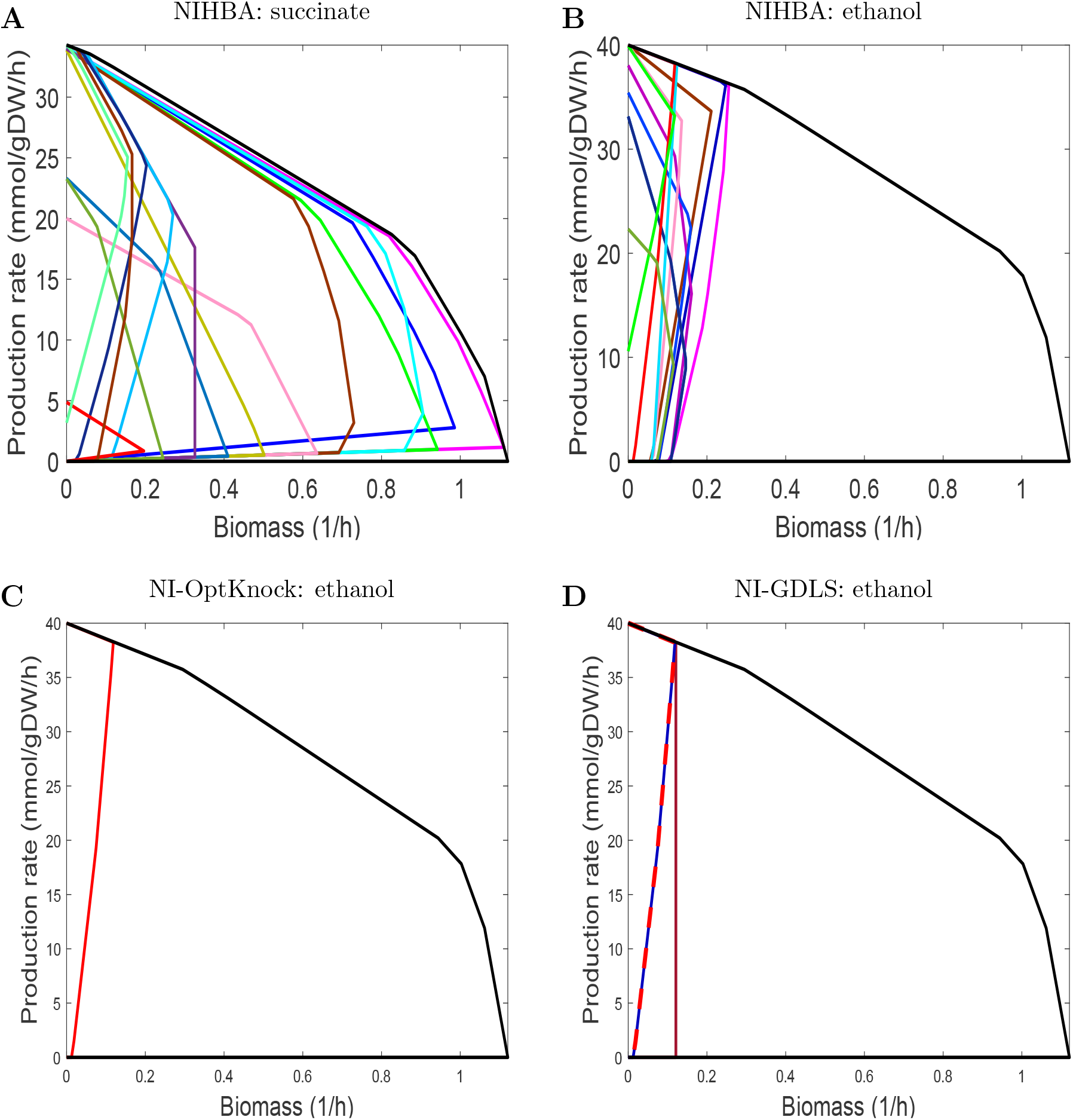
Production envelopes of diverse design solutions obtained by NIHBA for succinate and ethanol production. (**A**) Production envelopes of selected solutions for succinate production. (**B**) Production envelopes of selections solutions for ethanol production.

For ethanol, all the algorithms found feasible solutions with at most five knockouts, and all contain a solution with the maximum production rate, as illustrated in Fig. 4B–D (a small portion of solutions from NIHBA are displayed for readability). This shows that NIHBA has comparable performance in terms of optimality. Despite three solutions found from NI-GDLS, one of them is not growth coupled and the other two have the same production envelope, from which little can be gained about the tradeoff between cell growth and target production. In contrast, NIHBA found many solutions with diverse production envelopes, among which strong growth-coupled designs exist.

## 3 Discussion

The employment of bilevel optimisation for identifying genetic manipulations has been found very helpful for metabolic engineering. Most existing bilevel-based tools assume that cells always grow optimally, a biased optimisation principle that is found incorrect for mutants or certain microorganisms in some studies [Schuetz *et al.*, 2012; Segré *et al.*, 2002]. As a consequence, design strategies found by these tools may be biologically infeasible in spite of highest production rates at optimal growth. In addition, these tools involve solving a bilevel problem through big-M reformulation to a standard MILP that is suitable for commercial global search solvers like Gurobi and CPLEX. However, the resulting MILP is often large due to the genome scale of metabolic networks, and global search can be computationally prohibitive, particularly when a large design space (or numerous genetic manipulations) is allowed. Even if computationally tractable, the global search produces only a single solution. Furthermore, big-M formulation produces a weak MILP, leading to numeric issues such that no feasible solutions can be found. This paper have proposed to address biased assumptions from the point of view of game theory, leading to a network interdiction problem (NIP). The NIP is not handled using popular global search solvers, instead it is solved through an efficient hybrid Benders decomposition algorithm to lower computational costs and overcome numeric issues. The proposed approach, NIHBA, has shown its ability to obtain a large number of growth-coupled design strategies with diverse production phenotypes and achieve optimal production rates within an hour, regardless of the size of design space (the maximum allowable number of knockouts).

NIHBA uses a game theoretic framework to model the interaction (somehow competitive) between host cells and metabolic engineers. This eliminates the need for a biologically rigorous objective to reflect cells’ primary intentions, such as optimal growth or maximum energy generation [Schuetz *et al.*, 2012]. Therefore, design strategies found by NIHBA do not necessarily yield the best production at optimal growth. Instead, they guarantee non-zero production when cell growth is over a predefined survival threshold. NIHBA employs a hybrid Benders algorithm, HBA, to solve the NIP. Our case studies have demonstrated numerous advantages of this algorithm. First, it is free of numeric issues, making it much more stable than other MILP solvers, e.g. Gurobi and CPLEX. Second, it can be considered a parameter-free algorithm as opposed to other methods like GDLS that requires a setting of multiple parameters, although NIHBA uses a parameter *µ* for identifying Pareto optimal cuts. In practice, NIHBA is not sensitive until a value of *µ* < 1*e−*6 is used. Third, it is computationally efficient such that one hour on average is sufficient for NIHBA to identify high production solutions, and the runtime for a high production rate does not scale with the number of knockouts, which is not the case for existing methods. Last, it obtains numerous growth-coupled solutions in a single run. This is important as it not only helps understand the trade-off between target production and cell growth, but also provides the possibility to examine and test multiple solutions, from which the most promising design can be chosen for experimental implementation.

The proposed HBA is not limited to network interdiction problems. It can be applied to any bilevel or single-level optimisation problems that have complicating mixed-integer variables. Although promising, HBA needs improvements on convergence at late stages for optimality proof. Like other global search, an appropriate optimality gap or time limit may alleviate excessive exploration but cannot determine the optimality of solutions. Further improvements can be made along this direction to enhance the convergence of HBA. It is also noteworthy that multiple solutions by HBA are not searched in a systematic way. They may not form a perfect representative of the tradeoff between target production and cell growth. Therefore, more investigations are required to extract limited but well-diversified solutions in the search process of HBA.

Despite numerous solutions found by NIHBA, the selection of promising solutions poses a new challenge to decision makers. It is therefore important to have a good solution ranking approach. Solutions may be roughly ranked according to the frequency of individual knockouts in addition to their subsystem distribution or by a scoring system with manual settings [Schneider and Klamt, 2019]. However, a more systematic solution ranking is desirable, and this is left in our future work.

## 4 Materials and Methods

### 4.1 Flux balance analysis

A metabolic network of *m* metabolites and *n* reactions has a stoichiometric matrix *S* that is formed by stoichiometric coefficients of the reactions. Let *J* be a set of *n* reactions and *v*_*j*_ the reaction rate of *j* ∈ *J*, *Sv* represents the concentration change rates of the *m* metabolites. FBA aims at optimising a linear biological objective *c*^*T*^ *v* when the system is at steady state (i.e, the concentration change rate is zero for all the metabolites) and *v* is subject to thermodynamic constraints:

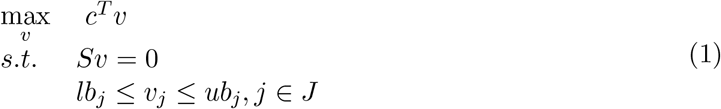

where *lb*_*j*_ and *ub*_*j*_ are the lower and upper flux bounds of reaction *j*, respectively. *c* is a weight vector specifying the degree of importance to the biological objective.

### 4.2 Network interdiction based strain design and reformulation

Network interdiction for strain design considers metabolic engineers as interdictors or adversaries who attempt to maximally disrupt host cells’ activity that biochemicals of interest are not overproduced due to homeostasis. The strain design task can therefore be formulated as a max-min problem:

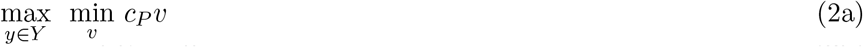

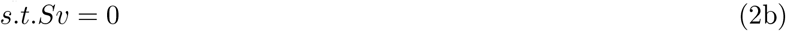

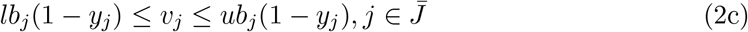

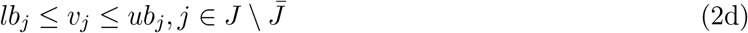

where *cP* is a coefficient vector for the target biochemical. 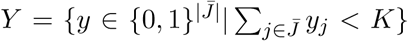 (*K* is the maximum allowable number of knockouts), and *y*_*j*_ indicates the reaction *j* is inactive (*v*_*j*_ = 0) if *y*_*j*_ = 1 and active otherwise. 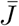 is a subset of *J*, containing candidate knockout reactions.

Observing that in the follower problem |*v*_*j*_|*y*_*j*_ = 0 always holds for all *j* ∈ *J*, we can eliminate all the flux constraints imposed by *y*_*j*_, i.e. eq.(2c), by rephrasing the inner objective function in a Lagrangian manner:

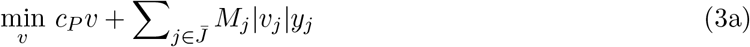

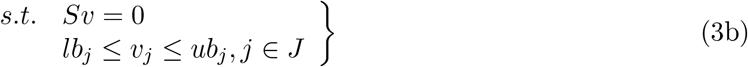

where *M*_*j*_ is a large positive Lagrange multiplier and *M* = (*M*_1_, …, *M*_|*¯J*|_). The reformulated follower problem is equivalent to the original problem in the sense that they have the same optimal value provided that *M*_*j*_ is sufficiently large for all *J* ∈ *¯J* such that *v*_*j*_ = 0 when *y*_*j*_ = 1. The value of *M*_*j*_ used in this work is around 100 (e.g. randomly drawn from [90,110]).

The reformulated follower function (3a) can be linearised by adding auxiliary variables *u*_*j*_ = *max*(*v*_*j*_, −*v*_*j*_). As a result, we have the reformulated bi-level framework:

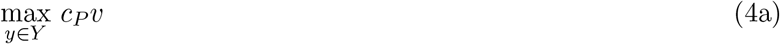

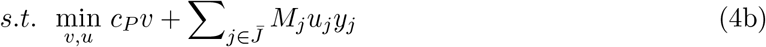

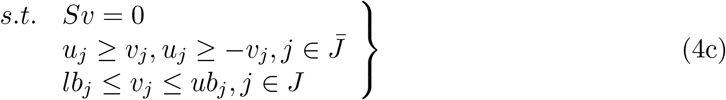

### 4.3 Hybrid Benders Algorithm

The bi-level problem (4) is reformulated to a standard MILP by applying LP duality to the follower problem (4b)–(4c). For simplicity, the resulting MILP is written in the following compact form

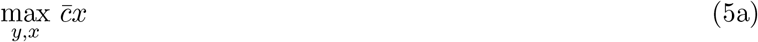

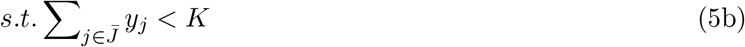

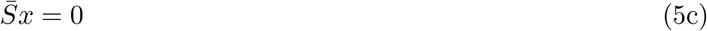

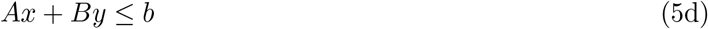

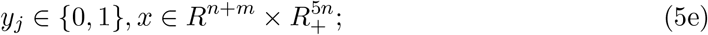

where 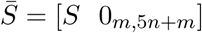, and

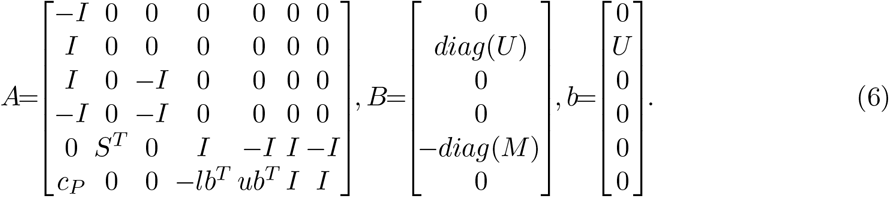

where *U* is a vector of maximum absolute flux for each reaction, i.e., *U*_*j*_ = *max*(|*lb*_*j*_|, |*ub*_*j*_|), ∀*j* ∈ *J*.

The single-level reformulation (5) can be solved, like OptKnock, by modern MILP solvers. However, the big-M terms in (5) lead to a week LP relaxation [Codato and Fischetti, 2006], therefore causing difficulties for MILP solvers. Besides, the model size of (6) increases rapidly for large metabolic networks, and as a result, a large-scale MILP has to be solved.

Benders decomposition avoids these drawbacks as it can deal with complicating binary variables and easy continuous variables separately. Like Benders decomposition [Codato and Fischetti, 2006], our hybrid Benders algorithm (HBA) decomposes (5) into a BIP master problem (MP) (7) and a LP slave problem (SP) (8) for fixed 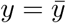:

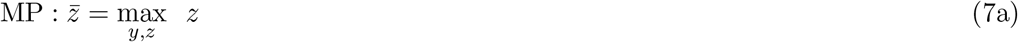

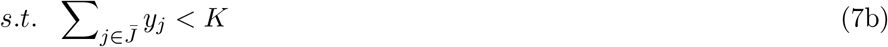

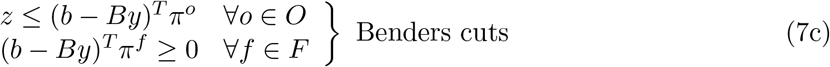

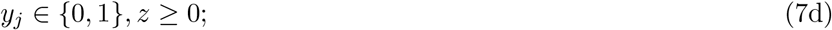

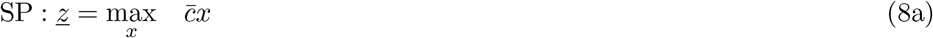

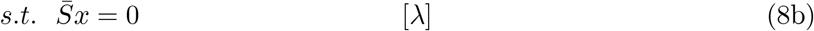

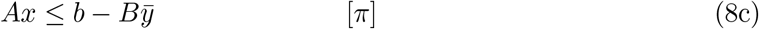

where *O* and *F* are sets that correspond to the extreme points *π*^*o*^ and extreme rays *π*^*f*^ of the dual of SP, respectively. In each iteration, the Benders decomposition algorithm derives the dual vector *π* from the SP (8) for 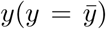 which is the solution to the MP in the previous iteration. In practice, a Benders cut is obtained by solving the dual of (8) rather than the primal. Two scenarios exist when solving the dual of (8): if the optimal value of the dual of SP is bounded, it means the SP is feasible, then an optimality cut *z* ≤ (*b* − *By*)^*T*^ *π*^*o*^ generated from the extreme point *π*^*o*^ is added to the MP; if it is unbounded, it means the SP is infeasible, then a feasibility cut (*b* − *By*)^*T*^ *π*^*f*^ ≥ 0 generated from the extreme ray *π*^*f*^ is added to the MP to avoid unboundedness of the dual of SP in future iterations.

The classic Benders decomposition is not able to generate effective Benders cuts rapidly for our strain design problem, and therefore requires a huge of iterations (consequently long computation time) before it converges. Here, we introduce a hybrid Benders algorithm (HBA) with two strategies to speed up the convergence process.

Figure 5 shows a simplified flowchart of HBA. An implementation of the algorithm in MATLAB can be found in https://sites.google.com/view/shouyongjiang/resources.

**Figure 5:**
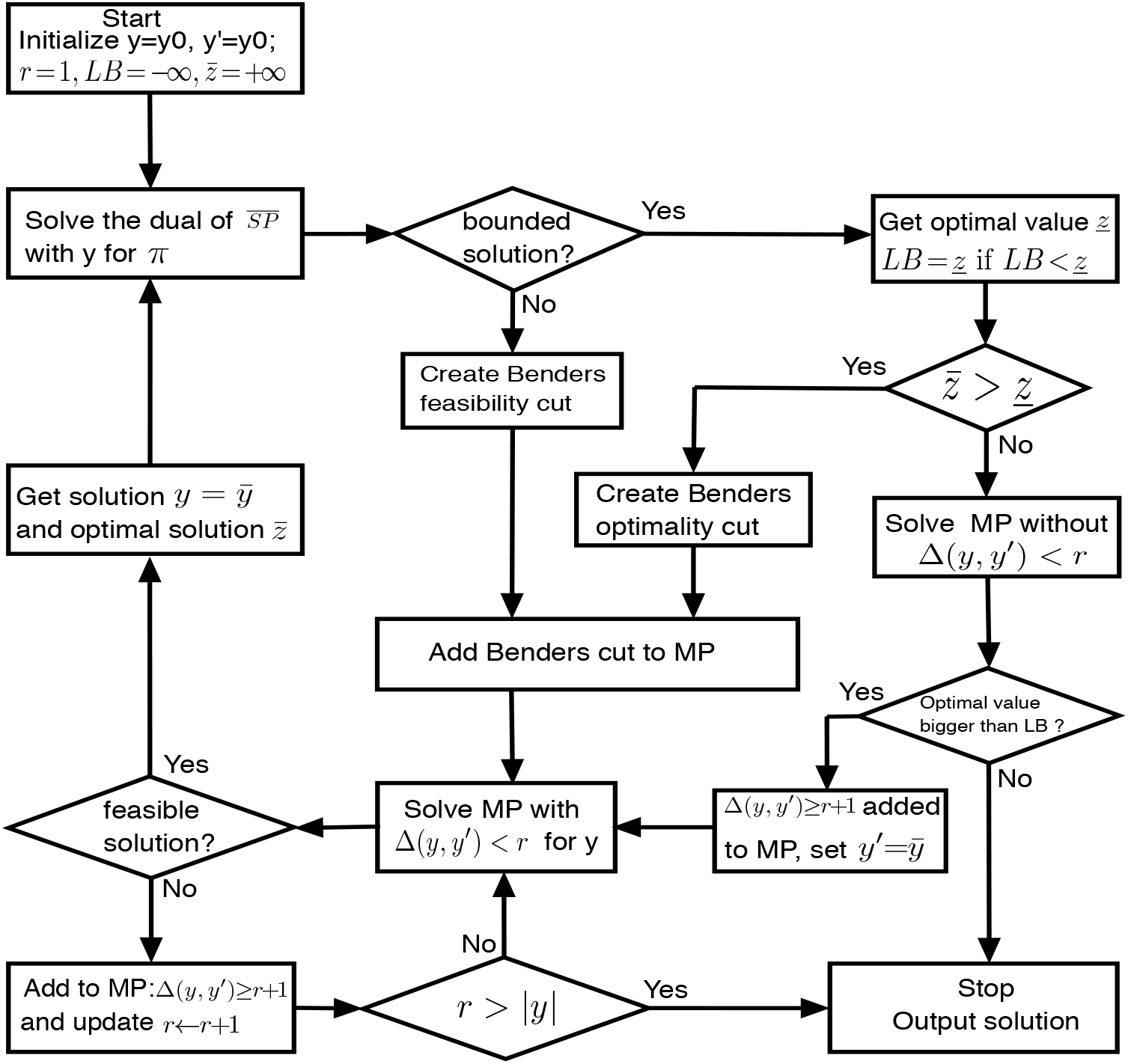
Flowchart of HBA.

#### 4.3.1 Pareto optimal cuts

Let *π*^*o*^ be the dual vector of *π* corresponding to (8), a standard Benders optimality cut is:

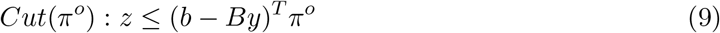

Since *π*^*o*^ may not be unique, it is important to select an effective cut *Cut*(*π*^*o*^). Magnanti and Wong [1981] proposed to use Pareto optimal cuts to improve convergence. *Cut*(*π*^*o*^) is said to be Pareto optimal if no other 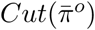 exists such that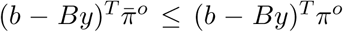 for any *y* ∈ *Y* and at least one *y* ∈ *Y* enables a strict inequality. There are a few methods available for identifying a Pareto optimal cut, but most of them have to solve the slave problem (8) twice, which may increase computational time significantly. We turn to the approach of Sherali and Lunday [2013] where a Pareto optimal cut can be generated by solving only once in each iteration a slightly different slave problem:

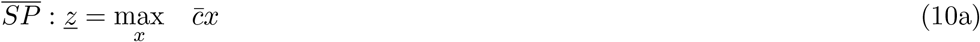

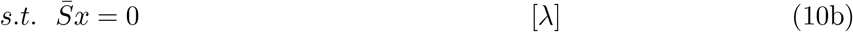

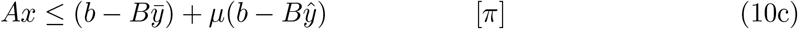

where *µ* is a sufficiently positive value and *ŷ* is a core point in the relative interior of the convex hull of *Y*. In this paper, *ŷ* is updated by 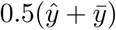 whenever a new feasible 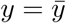 is produced in the iteration of Benders decomposition. *µ* is not calculated as in Sherali and Lunday [2013] but rather fixed to 1e-8 after multiple trials.

#### 4.3.2 Local branching

Another technique we used for accelerating the convergence of Benders decomposition is local branching, which is particularly effective when problems have binary variables [Baena *et al.*, 2018; Rei *et al.*, 2009]. Suppose *y′* is a feasible solution in *Y*, the idea behind local branching is to divide the feasible region of (7) into two subregions by the Hamming distance between *y* and *y′*:

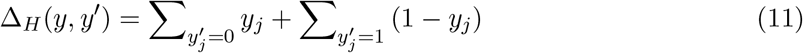

In every iteration of Benders decomposition, the master problem (7) is solved in the subregion ∆_*H*_(*y*, *y′*) < *r* (where *r* is a positive integer and the maximum is the cardinality *|y|* of *y*). This leads to two scenarios: there is either a feasible or infeasible *y* in the subregion ∆_*H*_ (*y*, *y′*) < *r*. If a feasible solution 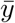 is obtained, Benders cuts are generated by solving (10) with 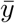. If not, it means the value of *r* may be too small, and ∆_*H*_(*y*, *y′*) > *r* + 1 is added to the master problem (7) to stop re-exploration in the neighbourhood of *y′* with the radius *r*. *r* is then increased by one at a time until ∆_*H*_(*y*, *y′*) < *r* renders the master problem (7) feasible. Note that *y′* has to be updated by 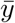 if 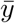 gives (7) an objective value worse than that of the slave problem (8), implying that no better solution can be obtained from the neighbourhood of *y′*.

### 4.4 Additional improvement strategies

HBA involves solving the MP (7) and SP (8) in a repeated manner. For efficiency, the following two strategies are used:

- Terminating the MP program prior to optimality. Suboptimal solutions to the MP are sufficient to generate valid Benders cuts. Therefore, the MP is terminated when a MIP Gap of 1 + 300/(*iter*^0.5^ + 1) (where *iter* is the iteration counter) is reached.
- Reversing local branching whenever the 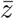 value of the MP is worse than <u>*z*</u> value of the SP. 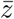 estimates the upper bound of the problem (2). 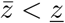 indicates the global optimum does not exist in the corresponding local branching and a reverse local branching should therefore be used.

### 4.5 Model reduction and candidate selection

The truncation of model size and candidate knockout set has great computational benefits. Genome-scale metabolic (GEM) models can be significantly simplified by compressing linear reactions and removing dead end reactions (those carrying zero fluxes). Likewise, many reactions can be excluded from consideration with *a priori* knowledge that, for example, they are vital for cell growth or their knockout is not likely to improve target production. We followed the model reduction procedure by Lun *et al.* [2009] and candidate selection procedure by Feist *et al.* [2010], resulting in a candidate set of 150*∼*350 reactions for different target products from the latest E. coli GEM iML1515 [Monk *et al.*, 2017] where the maximum uptake rates for glucose and oxygen are all 20 mmol/gDW/h.

### 4.6 Computational implementation

First of all, all the NI models were transformed into MILPs using duality theory [Burgard et al., 2003]. Then, the resulting MILPs were implemented in MATLAB 2018b to be compatible with the Cobra Toolbox 3.0 [Heirendt *et al.*, 2018] where we carried out simulations. All the MILPs were solved by Gurobi 7.5 [Gurobi Optimization, 2018] with both Heuristics and MIPFocus were set to 1 as suggested by Egen and Lun [2012]. A time limit of 2 hours was applied to each MILP while performing computations on Ubuntu 16.04 LTS with an Intel® CoreTM i5 Quad Core processor.

## Data and software availability

The data and software used and the tool developed are all available online:

- GEM model: iML1515 from BIGG database (bigg.ucsd.edu)
- Simulation software: Cobra toolbox 3.0 (https://opencobra.github.io/)
- Global solver: http://www.gurobi.com/
- NIHBA: https://sites.google.com/view/shouyongjiang/resources

## Acknowledgements

This work has been supported by the Engineering and Physical Sciences Research Council (EPSRC) for funding project “Synthetic Portabolomics: Leading the way at the crossroads of the Digital and the Bio Economies (EP/N031962/1)”.

## Author contributions

SJ and NK conceived the project. SJ developed the computational method and performed the computational simulations and analysis of the results. SJ wrote the draft of the paper. All authors read, edited, and approved the final manuscript.

